# IFN-*γ* signaling is required for the efficient replication of murine hepatitis virus (MHV) strain JHM in the brains of infected mice

**DOI:** 10.1101/2025.01.01.631031

**Authors:** Catherine M. Kerr, Macie A. Proctor-Roser, Srivatsan Parthasarathy, Joseph J. O’Connor, Jessica J. Pfannenstiel, Robin C. Orozco, Anthony R. Fehr

## Abstract

Neurotropic viruses are a major public health concern as they can cause encephalitis and other severe brain diseases. Many of these viruses, including flaviviruses, herpesviruses, rhabdoviruses and alphaviruses enter the brain through the olfactory neuroepithelium (ONE) in the olfactory bulbs (OB). Due to the low percentage of encephalitis that occurs following these infections, it’s thought that the OBs have specialized innate immune responses to eliminate viruses. Murine hepatitis virus strain JHM (JHMV) is a model coronavirus that causes severe encephalitis in mice and can access the brain through olfactory sensory neurons. We’ve shown that a JHMV Mac1-mutant virus, N1347A, has decreased replication and disease in the brains of mice. Here we further show that this virus replicates poorly in the OB. However, it is unknown which innate immune factors restrict N1347A replication in the OB. RNA seq analysis of infected olfactory bulbs showed that IFNγ was upregulated in the OB while IFN-*β* was barely detectable at 5 days post-infection. To determine if IFN-γ restricts JHMV N1347A replication, we utilized IFN-γ and IFN-γ receptor (IFN-γR) knockout (KO) mice. Surprisingly we found that JHMV WT and N1347A replicated very poorly in the OB and whole brains of both IFN-γ and IFN-γR KO mice following intranasal infection, though survival and weight loss were unaltered. Furthermore, we determined that microglia were the primary cells producing IFN-γ during the early stages of this infection. We conclude that IFN-γ is required for the efficient replication of JHMV in the brains of infected mice.

## INTRODUCTION

Viruses that infect the brain are a major public health threat, as they can lead to severe disease such as encephalitis, encephalomyelitis, flaccid paralysis, and aseptic meningitis. Neurotropic viruses can enter the brain through several mechanisms, including anterograde spread through sensory nerves, across the blood-brain barrier (BBB) as free virions, or through infection of immune cells that then enter the brain through the BBB. However, the most common route is through the olfactory bulb (OB) [1, 2]. Viruses that utilize OBs as an entrance to the central nervous system (CNS) are influenza A virus, herpesviruses, poliovirus, rabies virus, adenoviruses, Japanese encephalitis virus, West Nile virus, chikungunya virus, severe acute respiratory syndrome coronavirus 2 (SARS-CoV-2), and many others [2]. Importantly, SARS-CoV-2 utilizes the olfactory epithelium as a major entry point to the brain which can lead to encephalopathy and other neurological sequelae in some patients [3–8]. Despite the potential for severe consequences of a brain infection, severe disease following infection is somewhat rare, indicating that the cells in the OB have a robust innate immune response to limit virus replication and pathogenesis. However, we lack a full understanding of the main factors that limit virus replication in the olfactory bulb. Thus, understanding how cells in the olfactory bulb, including the olfactory neurons and epithelium respond to virus infection will be critical to develop novel methods to control neurotropic viruses.

The olfactory epithelium is adjacent to the respiratory epithelium in the upper airways where olfactory sensory neurons (OSNs) detect odorants which allow mammals to detect different scents. Axons of the OSNs interact with the dendrites of projection neurons in the glomeruli of the OB. Projection neurons send signals deeper into the brain, which are regulated by interneurons of the OB. Neurotropic strains of murine CoVs, specifically mouse hepatitis virus (MHV) strain JHM (JHMV) and its attenuated variant J2.2, cause acute encephalitis and CNS demyelination, respectively [9]. These viruses directly infect OSNs and enter the OB through anterograde transport via the olfactory nerve. JHMV then spreads *trans*-neuronally to connections of the OB which then allows it to traverse through the brain, ultimately leading to severe encephalitis or CNS demyelination [10, 11].

Coronavirus (CoVs) are positive-sense RNA viruses with a large genome of around 30kb. They have sixteen non-structural proteins (nsps) that are important for virus replication. The largest of these proteins is nsp3, which contains several domains, including ubiquitin-like domains, one or two papain-like protease domains, a deubiquitinase domain, a CoV-Y domain, and one or more macrodomains [12]. Of these macrodomains, the most N-terminal macrodomain, termed Mac1, is a conserved throughout the *Coronaviridae* family, and is also found in the *Togaviriadae, Hepeviridae*, and *Matonaviridae* families [13, 14]. Mac1 acts as an ADP-ribosylhydrolase, which removes ADP-ribose from target proteins [15]. We and others have used reverse genetics to define the role of Mac1 during CoV infection [16–26]. The primary Mac1 modification used in these studies is the mutation of a highly conserved asparagine to alanine (N-A), which is known to ablate the enzymatic activity of this protein [21, 25, 27]. These studies have found that recombinant viruses with this mutation replicate poorly and cause minimal disease in mice [18, 20, 21, 26, 27]. Specifically, the JHMV N1347A mutant virus replicates normally in standard cell lines used for JHMV but is highly attenuated in primary bone-marrow derived macrophages (BMDMs) and in mice [20, 28, 29]. Importantly, this attenuation is mediated at least in part by IFN-I, as N1347A replication and pathogenesis was mostly rescued in IFNAR^-/-^ cells and mice [28]. In addition, JHMV N1347A was more sensitive to pre-treatment of BMDMs with IFN-*β* than WT virus. Finally, microglia were critical for the restriction of N1347A in mice, as mice depleted of microglia were highly sensitive to N1347A, unlike mice with intact microglia, indicating that microglia are the likely producers of IFN-I during JHMV infection [11].

ADP-ribosylation is a common post-translational modification of target proteins. It is catalyzed by ADP-ribosyltransferases, the most common of which are PARPs, which can add either a single ADP-ribose moiety (MARylation) or a chain of moieties (PARylation) [30]. Several PARPs are also interferon stimulating genes (ISGs), as they are unregulated upon IFN stimulation [28, 31, 32]. As such, a number of PARPs have antiviral activity [33–40]. Previously, using pan-PARP inhibitors we demonstrated that PARPs inhibit JHMV replication in the absence of Mac1 activity [28]. More recently, we showed that PARP12, an ISG, is required to for the restriction of N1347A replication in cell culture and *in vivo* in a cell and tissue specific manner [41]. JHMV N1347A replication was increased to WT virus levels in PARP12*^-/-^* BMDMs. However, in mice the absence of PARP12 resulted in several different levels of virus restoration depending on the tissue and route of infection. MHV-A59 N1348A (same mutation as in JHMV) replication was increased to WT levels in the livers of PARP12^-/-^ mice compared to PARP12^+/+^ mice and the pathology in the liver was partially restored. Following an intracranial infection of PARP12^-/-^ mice, JHMV N1347A replication and lethality was increased compared to PARP12^+/+^ mice but not to WT virus levels. Finally, we found no difference in N1347A replication or virulence following an intranasal infection, which enters the brain through OSNs, of PARP12^-/-^ mice compared to WT mice [41]. This indicates that PARPs, specifically PARP12, are involved in restricting Mac1 mutant virus replication *in vivo* but that there must be additional immune factors in the OBs that control N1347A replication and its disease outcomes.

In this study, we aimed to identify host innate immune factors, specifically PARPs, that might restrict N1347A in the OBs of mice. We found that IFN-*γ*, but not IFN-I, was highly upregulated in OBs early after infection, but surprisingly found that, unlike IFN-I, IFN-*γ* was required for the efficient replication of JHMV in mice, as JHMV WT and N1347A replication was dramatically reduced in IFN-*γ*^-/-^ and IFN-*γ*R^-/-^ mice. These results provide new insight into the impact of IFN-*γ* on the replication of a neurotropic CoV.

## RESULTS

### JHMV-N1347A replicates poorly in the olfactory bulbs following intranasal infection

We recently demonstrated that JHMV N1347A replication is increased following an intracranial infection of PARP12^-/-^ mice compared to WT mice, but not following an intranasal infection [41]. Therefore, we tested the ability of N1347A to replicate at the initial site of an intranasal infection, the OBs. WT (+/+) mice were infected with either the WT virus or N1347A virus, then OBs were harvested at peak titer (5 days post-infection (dpi)) to determine the levels of virus replication. We could not identify significant levels of infectious virus prior to this time point in the olfactory bulb (data not shown). We found that levels of N1347A infectious virus and genomic RNA levels were significantly reduced compared to the WT virus in the OB **(Fig. 1A-B)**. Additionally, less N protein was observed by immunohistochemistry in N1347A infected olfactory bulbs compared to WT infected olfactory bulbs **(Fig. 1C)**. Taken together, this data indicates that N1347A replicates poorly in the OB and is likely restricted by one or more unknown host factors.

**Fig 1.**
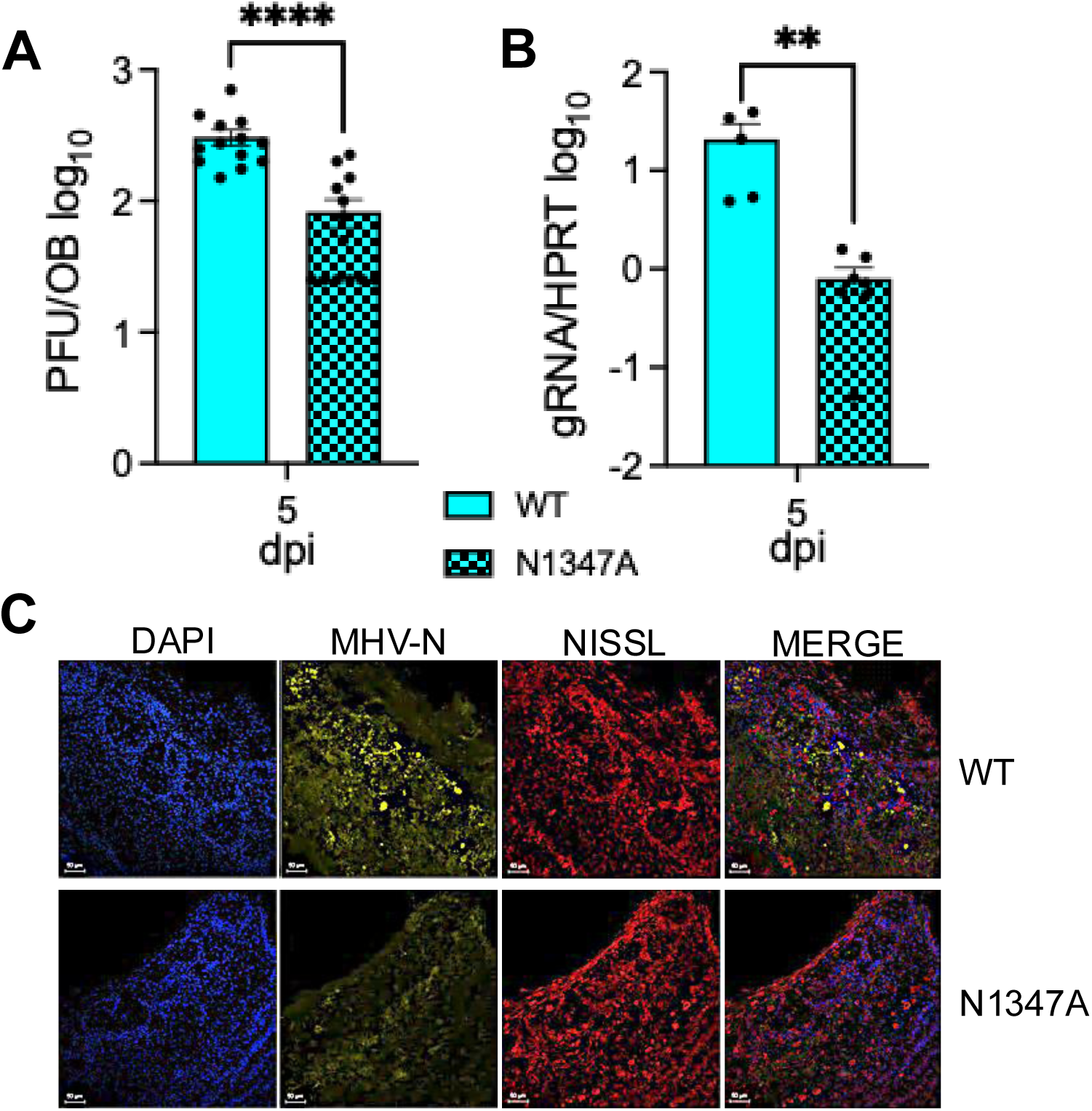
JHMV N1347A has decreased replication in the olfactory bulbs of mice. (A-C) C57BL/6 mice were infected with infected i.n. with 10^4^ PFU of JHMV WT or N1347A. Brains were harvested at 5 dpi and viral load was determined via plaque assay (A), genomic RNA levels (gRNA) were determined by RT-qPCR (B) and olfactory bulb sections were stained for MHV nucleocapsid protein (yellow), NISSL (Neurotrace^TM^, red), and DAPI (blue) by IHC (C). The data in A is the combined results of 3 independent experiments. n=13 mice per group. The data in B is the combined result of 2 independent experiments. n=4 mice (WT) and n=6 mice (N1347A). The images in C are from 1 experiment representative of 2 independent experiments. Scale bar is 100 *μ*m. n=2-4 mice per group. Statistics were determined using a Mann-Whitney t-test.

### IFN-γ is upregulated in infected olfactory bulbs

Next, we utilized RNAseq to identify immune factors in the olfactory bulb that could be responsible for the observed reduction in N1347A replication. Interestingly, compared to naïve mice, we observed that IFN-II, or IFN-γ, but not IFN-I, was upregulated in the olfactory bulb upon an N1347A infection **(Fig. 2A-B and Table S1)**. In addition to IFN-γ, several other ISGs were upregulated as well, including ISG15, ISG20, CXCL9, CXCL10, CXCL11, IFIT2, IFIT3, RSAD1, RSAD2, and several PARPs **(Fig. 2A-B)**. We then used qPCR to confirm that IFN-γ, but not IFNβ, was upregulated in the OBs following JHMV infection **(Fig. 2C)**. We’ve recently shown that pre-treatment of cells with IFN-γ in cell culture induces PARP expression [18], so despite the lack of IFN-I, we analyzed the OBs for PARP expression. We identified 7 PARPs that were upregulated in OBs following JHMV infection, including PARP3, 7, 9, 11, 12, 13, and 14 **(Fig. 2D)**, one or more of which likely contribute to the repression of N1347A in OBs.

**Fig 2.**
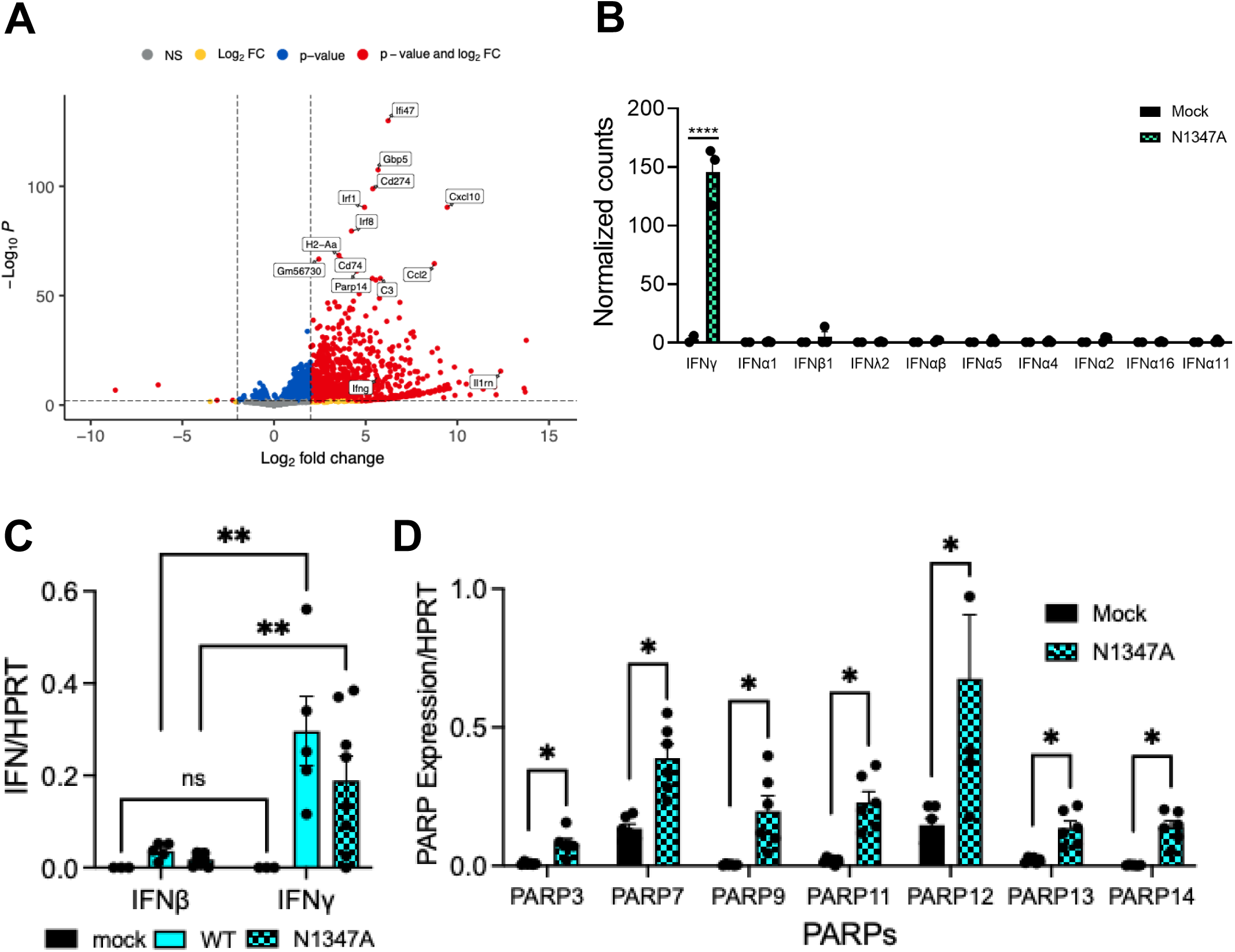
IFNγ and several PARP genes are upregulated in infected olfactory bulbs of JHMV infected mice. (A) C57BL/6 mice were either mock infected or infected i.n. with 1×10^4^ PFU N1347A virus. Olfactory bulbs were harvested at 5 dpi in Trizol, and total RNA was isolated. The total RNA from the samples were analyzed by RNAseq to determine the full transcriptome in the olfactory bulb following infection and is represented by a volcano plot indicating *<u>d</u>*ifferentially *<u>e</u>*xpressed *<u>g</u>*enes (DEGs) between mock and N1347A infected mice. Vertical dashed lines highlight absolute log2 fold-change (LFC) values > 2. Genes that are significant and have LFC values > 2 are shown in red, those that are significant but with LFC values < 2 are in blue, genes that are not significant but have LFC values > 2 are in yellow, and genes that are not significant and with LFC values < 2 are in grey. n=2 mice (mock); n=3 mice (N1347A). (B) Normalized counts of individual IFN genes between mock and N1347A infected mice from the RNAseq data show in 2A were calculated and represented in form of a bar graph. (C) C57BL/6 mice were either mock infected or infected i.n. with 1×10^4^ PFU WT or N1347A virus. Olfactory bulbs were harvested at 5 dpi in Trizol, and total RNA was isolated. The levels of IFN*β* and IFN*γ* mRNA transcripts were analyzed by qRT-PCR normalized to HPRT. n=3 mice (mock); n=5 mice (WT); n=8 mice (N1347A). Statistics were determined using a 2-way ANOVA. (D) C57BL/6 mice were either mock infected or infected i.n. with 1×10^4^ PFU N1347A virus. Olfactory bulbs were harvested at 5 dpi in Trizol, and total RNA was isolated. The levels of PARP mRNA transcripts were analyzed by qRT-PCR normalized to HPRT. n=6 mice (mock); n=6 mice (N1347A). Statistics were determined using a Mann-Whitney t-test.

### IFN-γ is required for efficient replication of JHMV in the brain

Because we previously demonstrated that IFN-γ leads to upregulation of PARPs, we hypothesized that in the absence of IFN-γ or IFN-γ signaling JHMV N1347A replication and pathogenesis would be enhanced. However, what we observed was quite the opposite. At 5 dpi, JHMV WT viral loads were significantly reduced in both IFN-γ^-/-^ and IFN-γR^-/-^ OBs, and N1347A viral loads were extremely low in all mice, clearly demonstrating that deletion of IFN-γ or IFN-γ signaling did not enhance JHMV N1347A replication in the OBs **(Fig. 3A)**. We next analyzed viral loads in the whole brains of these mice at 5 dpi and found that both JHMV WT and N1347A viral loads were significantly decreased in IFN-γ^-/-^ and IFN-γR^-/-^ mice **(Fig. 3B)**. This indicates that IFN-γ may be required for efficient JHMV replication following intranasal inoculation. To test this hypothesis using a separate measure of virus replication, we measured viral genomic and sub-genomic RNA of WT virus in the olfactory bulbs **(Fig. 3C)** and whole brains **(Fig. 3D)** of WT, IFN-γ^-/-^, and IFN-γR^-/-^ mice. In most cases, viral RNA levels were reduced in IFN-γ^-/-^, and IFN-γR^-/-^ mice, with statistically significant reduction of viral RNA in the whole brains. To further demonstrate that viral replication is repressed in the olfactory bulb, we stained infected WT and IFN-γ^-/-^ olfactory bulbs for the CoV N protein using immunohistochemistry to identify cells with actively replicating virus. JHMV WT infected OBs from WT mice had robust N protein staining, while N1347A staining was reduced compared to WT virus **(Fig. 3E-F)**. However, the expression of N protein in both the OB and forebrain of IFN-γ^-/-^ mice was substantially reduced compared to WT mice following infection of both JHMV WT and N1347A **(Fig. 3E-F and Fig. S1)**. In combination, these results indicate that IFN-γ is required for efficient replication of JHMV in the brain following intranasal infection.

**Fig 3.**
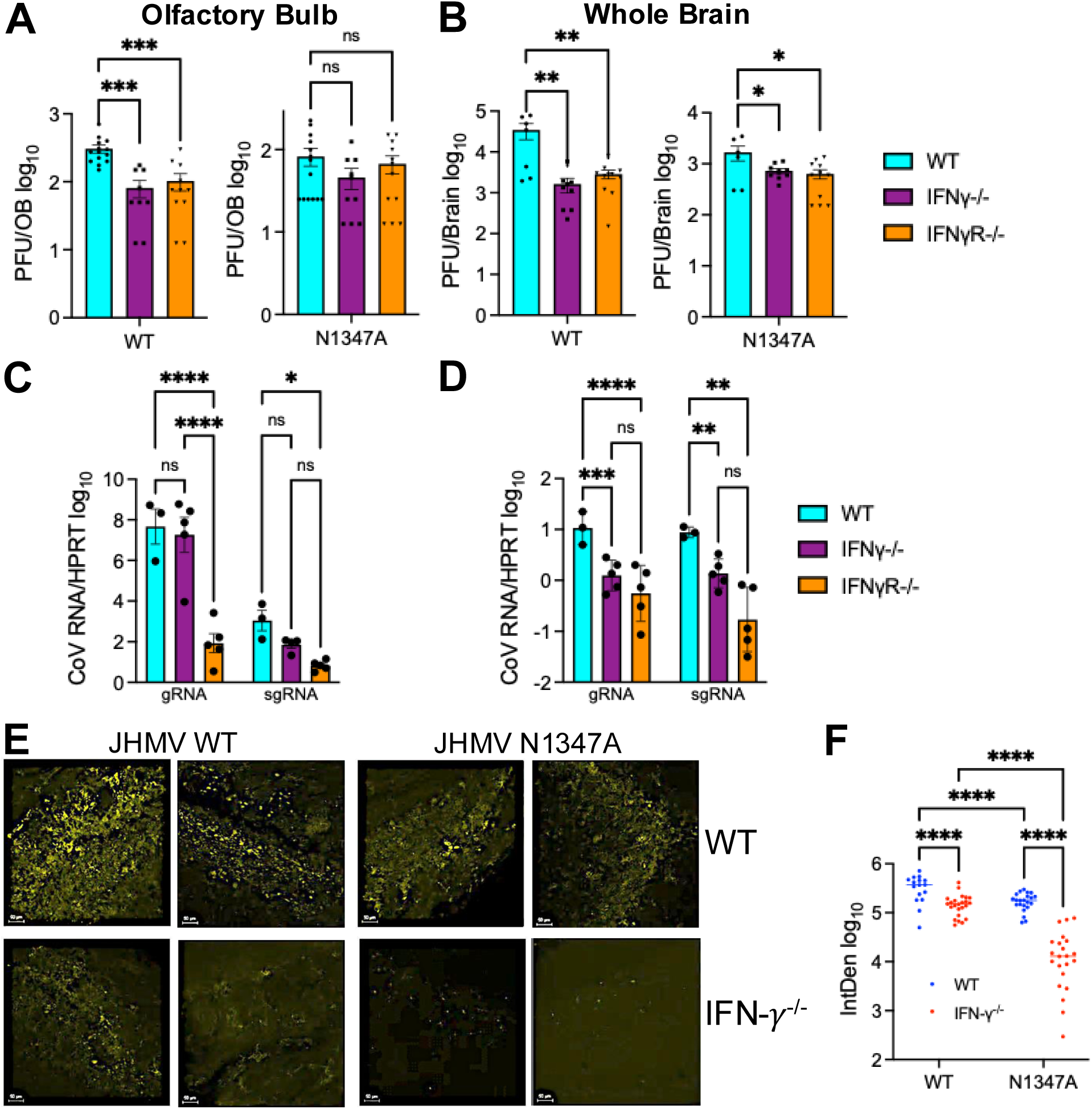
JHMV replicates poorly in the brains of IFN-*γ* and IFN*γ*R null mice. (A-B) C57BL/6 WT, IFN-*γ*^-/-^, and IFN-*γ*R^-/-^ mice were infected i.n. with 1×10^4^ PFU WT or N1347A virus. Olfactory bulbs (OB) (A) or whole brains (B) were harvested at 5 dpi and viral loads were measured by plaque assay. Data from WT mice in (A) is the same data in Fig. 1A. It is placed here as a comparison to the other genotypes. (A) n=13 (WT/WT), n=13 (WT/N1347A), n=8 (IFN-*γ*^-/-^ /WT), n=9 (IFN-*γ*^-/-^/N1347A), n=10 (IFN-*γ*R^-/-^/WT), n=11 (IFN-*γ*R^-/-^/N1347A). (B) n=6 (WT/WT), n=6 (WT/N1347A), n=8 (IFN-*γ*^-/-^/WT), n=9 (IFN-*γ*^-/-^/N1347A), n=10 (IFN-*γ*R^-/-^/WT), n=11 (IFN*γ*R^-/-^/N1347A). Statistics were determined by a one-way ANOVA. (C-D) Indicated mouse strains were infected i.n. with 1×10^4^ PFU WT virus. Olfactory bulbs (C) or whole brains (D) were harvested a 5 dpi and viral genomic and sub-genomic RNA was measured by RT-qPCR. n=3 (WT/WT), n=5 (IFN-*γ*^-/-^/WT), n=10 (IFN-*γ*R^-/-^/WT). Statistics were determined by a one-way ANOVA. (E) C57BL/6 WT, and IFN-*γ*^-/-^ cells were infected i.n. with 1×10^4^ PFU WT or N1347A virus. At 5 dpi olfactory bulbs were fixed and sections were stained for MHV nucleocapsid (N) protein (yellow) by IHC. Each image represents a different mouse. n=2 mice for each group. Each image represents a different mouse and WT images include images from the same mice as in Fig. 1. Co-staining of images with DAPI (nuclei) and Nissl (Neurotrace^TM^) as controls can be seen in Fig. S1. (F) Quantification of all images taken for each group. Each dot represents a different image. Statistics were determined using a 2-way ANOVA.

### The loss of IFN-γ or IFN-γ signaling does not impact JHMV-induced lethality

We next asked if the limited virus replication following JHMV intranasal infection in mice lacking IFN-γ or IFN-γ signaling would impact viral pathogenesis. WT, IFN-γ^-/-^, or IFN-γR^-/-^ mice were infected with WT JHMV and monitored for weight loss and lethality. Using a dose of 1×10^4^ PFU JHMV, we observed no difference in the lethality or weight loss in any of these mice **(Fig. 4A)**. We next tested if differences in pathogenesis might only be observed at lower doses of virus, so we infected mice with both 3×10^3^ and 1×10^3^ PFU. Despite these lower doses, we observed no consistent differences in the pathogenesis of JHMV in IFN-γ^-/-^ or IFN-γR^-/-^ mice **(Fig. 4B-C)**. We conclude that despite reduced levels of virus replication, the loss of IFN-γ or IFN-γ signaling does not affect the ability of JHMV to cause severe disease in mice. It also suggests that virus replication is not necessarily coupled to pathogenesis in the brains of mice infected with JHMV.

**Fig 4.**
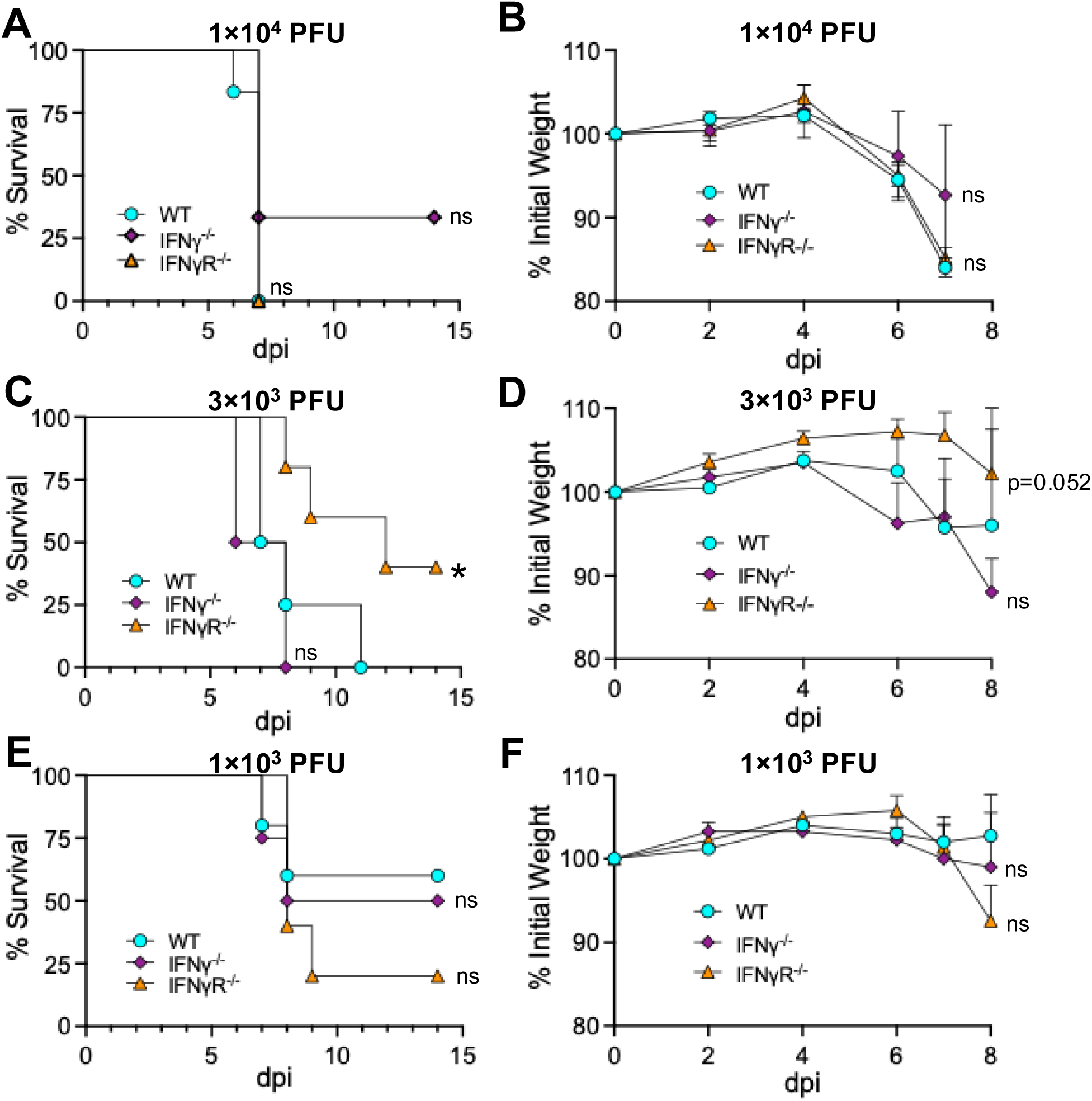
JHMV survival and weight loss is not altered IFN-*γ* null mice. (A-B) C57BL/6 WT, IFN-*γ*^-/-^, and IFN-*γ*R^-/-^ mice were infected i.n. with 1×10^4^ (A-B), 3×10^3^ (C-D), or 1×10^3^ (E-F) PFU WT virus. Mice were monitored daily for survival (A,C,E) and weight loss (B,D,F). (A-B) n=6 (WT), n=3 (IFN-*γ*^-/-^), n=7 (IFN-*γ*R^-/-^). (C-D) n=4 (WT), n=4 (IFN-*γ*^-/-^), n=5 (IFN-*γ*R^-/-^). (E-F) n=5 (WT), n=4 (IFN-*γ*^-/-^), n=5 (IFN-*γ*R^-/-^). Statistics were determined using a Kaplan-Meier survival analysis (A,C,E) or an ordinary one-way ANOVA (B,D,F).

### The loss of IFN-γ or IFN-γ signaling does not impact JHMV replication in ex vivo BMDMs

To test whether JHMV replicates poorly in IFN-γ^-/-^ or IFN-γR^-/-^ cells *ex vivo*, we harvested bone marrow from IFN-γ^-/-^ or IFN-γR^-/-^ mice and differentiated them into bone-marrow derived macrophages (BMDMs), which are susceptible to JHMV. BMDMs were infected at an MOI of 0.1 and harvested at 20 hpi and infectious virus and viral RNA were measured by plaque assay and qPCR, respectively. There were no significant differences in the production of infectious virus or viral RNA following JHMV WT or N1347A virus infection IFN-γ^-/-^ or IFN-γR^-/-^ cells when compared to WT cells **(Fig. 5A and Fig. S2)**. Of note, we have previously found that BMDMs produce very little IFN-γ (data not shown). These results indicates that cells lacking IFN-γ do not have an intrinsic defect in virus replication. Next, we pre-treated BMDMs with recombinant IFN-γ to test its ability to directly affect virus replication. The exogenous addition of IFN-γ to BMDMs reduced virus replication **(Fig. 5B)**. These results demonstrate that IFN-γ does not directly enhance virus replication and that the effects observed in the brain following intranasal infection are likely indirect effects of IFN-γ on cells in the brain.

**Fig 5.**
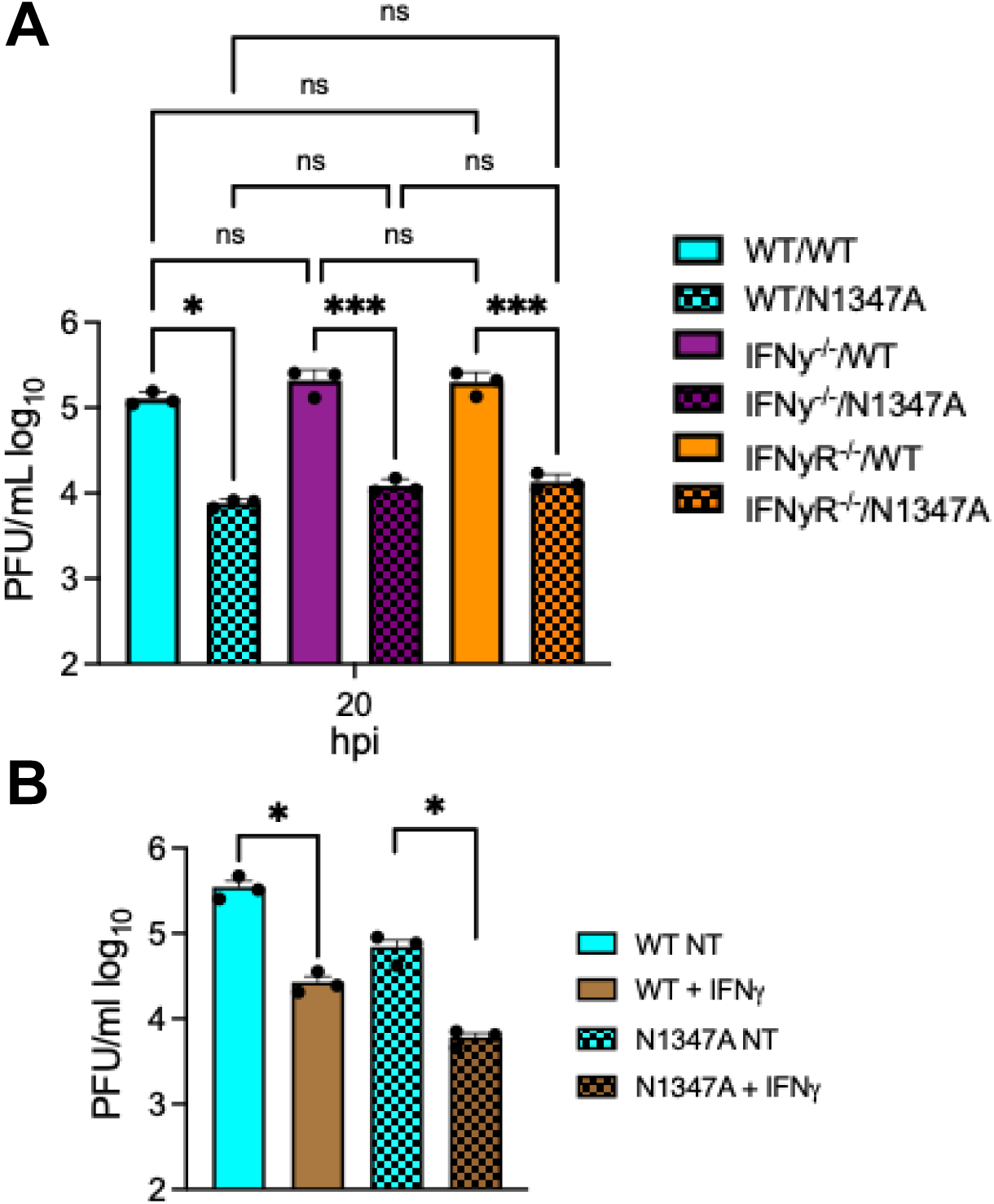
JHMV replication is unaffected by the lack of IFN-*γ* and is repressed by the addition of exogenous IFN-*γ* in bone-marrow derived macrophages. (A) C57BL/6 WT, IFN-*γ*^-/-^, and IFN-*γ*R^-/-^ bone-marrow derived macrophages (BMDMs) were harvested from mice and differentiated into M2 macrophages as described in Methods. BMDMs were infected with WT and N1347A virus at an MOI of 0.1 and cells and supernatants were collected at indicated time points and infectious virus was measured by plaque assay. (B) BMDMs were pretreated with IFN-*γ* for 20 hr and then infected with JHMV at an MOI of 0.1 and cells and supernatants were collected at indicated time points and infectious virus was measured by plaque assay. (A-B) are from one experiment representative of two independent experiments. Statistics in A-B were determined by an ordinary one-way ANOVA.

### The loss of IFN-γ signaling does not impact cytokine production or the immune cell profile of JHMV-infected brains

To try to explain how IFN-γ might alter virus replication *in vivo*, we next tested whether JHMV infection of IFN-γ^-/-^ or IFN-γR^-/-^ mice induce increased levels of IFN-I or other cytokines that could lead to reduced viral loads in the brain. To determine the levels of cytokines in the brain, we harvested both the olfactory bulbs and whole brains from JHMV WT infected mice at 5 dpi, prepared RNA, and measured the levels of IFN-*β*, IFN-*γ*, IL-1*β*, CXCL10, and IL-6 mRNA in WT, IFN-γ^-/-^ or IFN-γR^-/-^ mice using quantitative PCR **(Fig. 6A)**. The levels of most cytokines were not statistically different between the different strains of mice, except for CXCL-10, which was reduced in IFN-γ^-/-^ or IFN-γR^-/-^ mice. This is likely due to either the lower levels of virus replication or the lack of IFN-γ signaling, which is known to induce CXCL-10 **(Fig. 3C-D)**. We conclude that the lack of IFN-γ signaling does not result in broad changes in cytokine production following JHMV infection.

**Fig 6.**
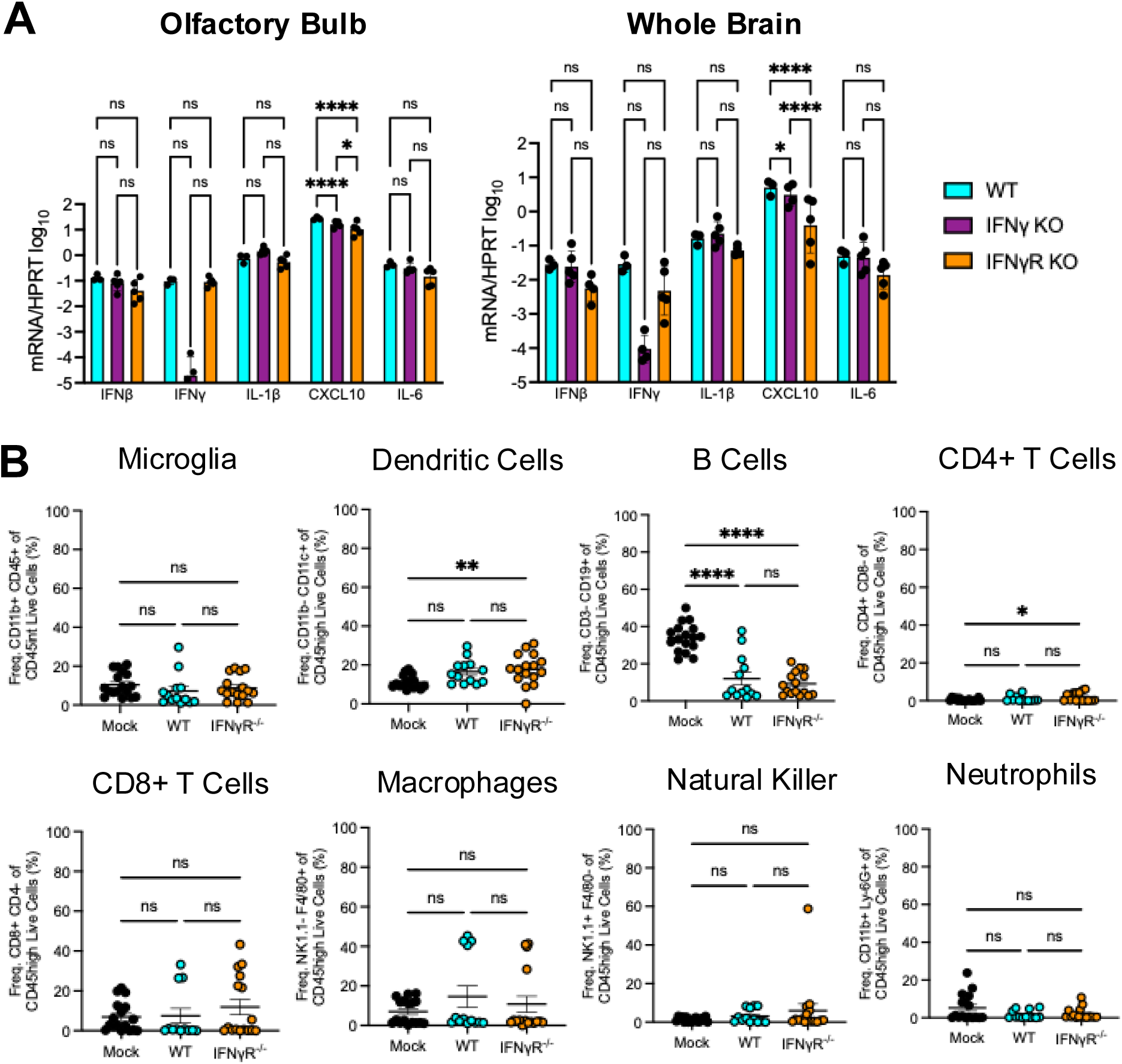
WT and IFN-*γ*R-/- have similar amounts of cytokines and frequencies of immune cells in the brain following infection. (A) C57BL/6 WT, IFN-*γ*^-/-^, or IFN-*γ*R^-/-^ mice were infected i.n. with 1×10^4^ PFU WT virus. Olfactory bulbs (left panel) and whole brains (right panel) were harvested at 5 dpi and total RNA was collected and cytokine mRNA was measured by RT-qPCR, normalized to HPRT. n=3 mice (WT); n=5 mice (IFN-*γ*^-/-^); n=5 mice (IFN-*γ*R^-/-^). Statistics were determined using an ordinary one-way ANOVA. (B) C57BL/6 WT or IFN*γ*R-/- mice were either mock infected or infected i.n. with 1×10^4^ PFU WT virus. Whole brains were harvested at 5 dpi and cells were stained for immune cell markers and analyzed by flow cytometry. n=17 mice (mock); n=13 mice (WT); n=16 mice (IFN-*γ*R-/-). Statistics were determined using an ordinary one-way ANOVA.

Next, we analyzed the immune cell profile in JHMV infected WT and IFN-γR^-/-^ mice by flow cytometry using naïve WT mice as controls. Again, we harvested the brains of infected mice at 5 dpi and measured the frequency of several innate and adaptive immune cells including microglia, dendritic cells (DCs), macrophages, natural killer cells (NK), neutrophils, B cells, and CD4+/CD8+ T cells **(Fig. 6B)**. We found no significant difference in any cell type between WT and IFN-γR^-/-^ infected mice.

### Microglia, CD4+ T cells, and Macrophages produce the highest amount of IFN-γ during JHMV infection

Following JHMV intranasal infection, we observed a stark increase in IFN-γ at 5 dpi **(Fig. 2)**, which prompted us to ask which cells were producing IFN-γ in these mice following infection and if the lack of IFN-γ receptor signaling impacts IFN-γ production. IFN-γ is typically produced by T cells or NK cells, however, this does not preclude other cells from producing IFN-γ during JHMV infection. To identify which cells produce IFN-γ, we harvested whole brains of JHMV infected WT and IFN-γR^-/-^ mice and measured IFN-γ production using intracellular flow cytometry. During JHMV infection in WT mice, there were no statistically significant differences detected between the frequency of IFN-γ + cell types, though microglia was the only cell type to produce IFN-γ in each mouse that was infected (**Figure 7A)**. However, in IFN-γR^-/-^ mice, there were significantly more IFN-γ + NK cells, though this difference was largely driven by two outliers **(Figure 7A)**. Using gMFI, microglia, CD4 T cells, and macrophages had the highest expression of IFN-γ in both infected WT and IFN-γR^-/-^ mice, while CD8+ T cells, NKs, and DCs produced substantially less IFN-γ **(Figure 7B)**. Furthermore, the MFI levels were nearly the same between the WT and IFN-γR^-/-^ mice indicating that IFN-γ receptor signaling does not alter IFN-γ production during JHMV infection. We conclude that microglia, CD4+ T cells, and macrophages contribute to the overall production of IFN-γ following an intranasal infection with JHMV.

**Fig 7.**
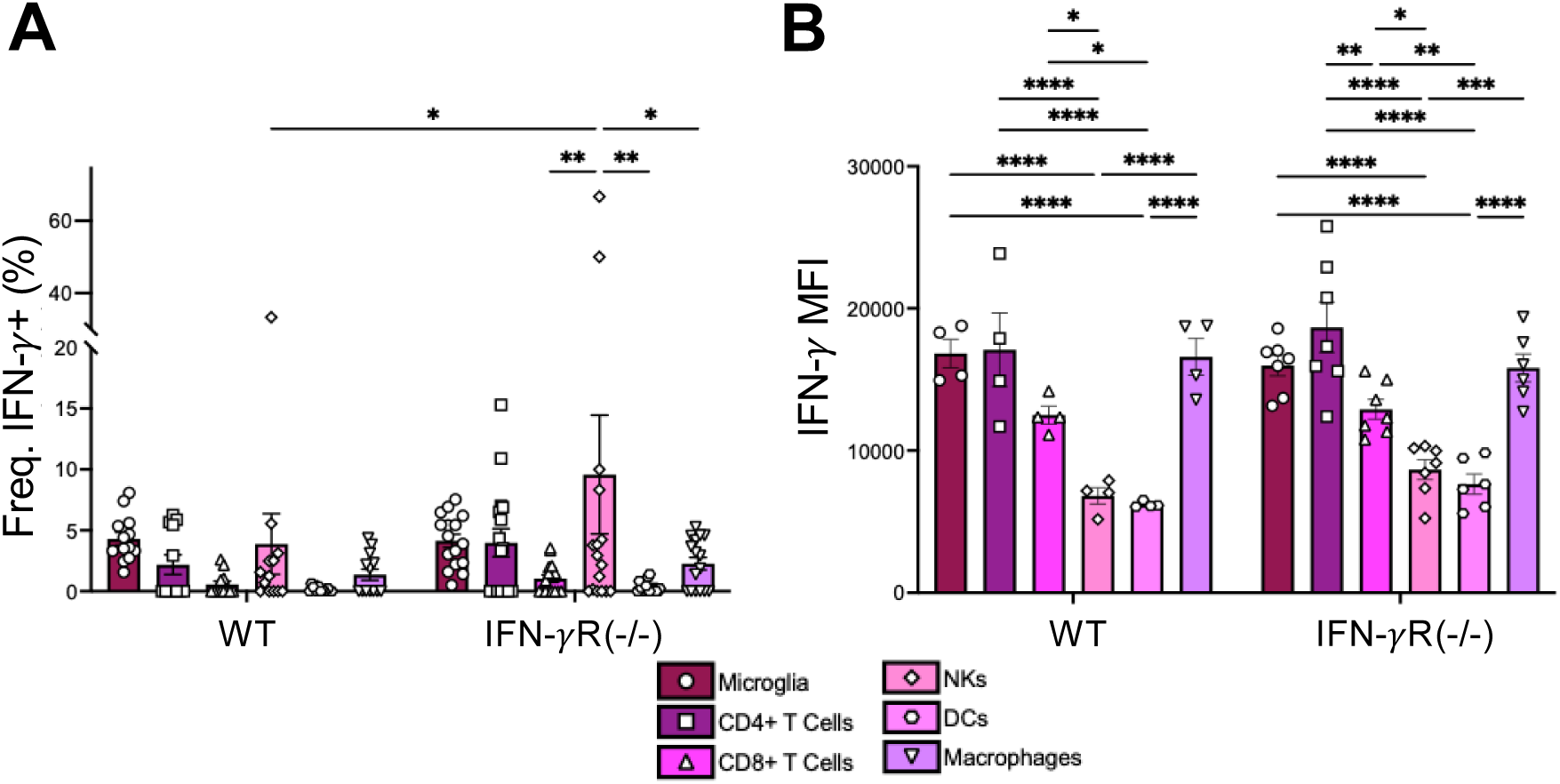
IFN-γ is produced by microglia, CD4+ T cells, and macrophages in the brains of JHMV infected mice. C57BL/6 WT or IFN-*γ*R-/- mice were either mock infected or infected i.n. with 1×10^4^ PFU WT virus. Whole brains were harvested at 5 dpi and IFN*γ* staining from each cell population was analyzed by flow cytometry. (A) The frequency of IFN-*γ*+ cells and (B) expression of IFN-*γ* (gMFI) for each cell type shown. Cell types analyzed were microglia (plum, circle), CD4+ T cells (magenta, square), CD8 T cells (bright pink, triangle), NK cells (carnation, diamond), DCs (pink, hexagon), and macrophages (purple, upside down triangle). Each point represents a mouse. The data in (A) is a pooled data from 3 independent studies while the data in (B) is from a representative experiment. Statistics were determined using a two-way ANOVA.

## DISCUSSION

In this study, we explored the role of IFN-γ in an intranasal infection of MHV-JHM, or JHMV. Our previous studies found that in WT mice a Mac1 mutant, N1347A, had a severe replication defect and caused minimal disease following i.n. infection of the brain [20, 28]. While Mac1-mutant virus had enhanced replication and led to increased disease in PARP12^-/-^ mice following both i.c. and i.p. infections, Mac1-mutant replication or disease was not enhanced at all in PARP12^-/-^ mice following an i.n. infection [41]. Unlike an i.c. infection, the olfactory bulb is the initial site infected following i.n. injection [10]. Thus, we hypothesized that there must be another PARP or innate immune factor present in the olfactory bulbs that restricts the replication of N1347A. Using RNA seq, we showed that IFN-γ, but ironically not IFN-I, was upregulated in the olfactory bulbs of mice following JHMV infection.

IFN-γ is the only known type II interferon. It is a component of both the innate and adaptive immune responses and is primarily produced by activated natural killer (NK) cells, CD4^+^ T helper-1 cells (T_H_1), and CD8^+^ cytotoxic T cells [42]. Specific functions of IFN-γ include T-cell activation, DC maturation, and polarization of macrophages to the M1 phenotype [43]. IFN-γ also has antiviral properties. For example, during a herpes simplex virus 2 (HSV-2) infection, the absence of IFN-γ increased virus replication and decreased survival of infected mice [44]. IFN-γ has also been shown to have a protective effect in mice against a challenge of influenza A virus (H1N1) [45]. This study found that WT mice were able to clear a viral challenge of H1N1 following an immunization with H3N2, while IFN-γ^-/-^ mice were not. The increased viral loads of H1N1 in IFN-γ^-/-^ mice indicate the importance of IFN-γ to clear subsequent virus infections [45]. It was recently shown that the administration of IFN-γ prior to infection provides protection to mice from a SARS-CoV-2 infection, and also suggested that the presence of IFN-γ likely decreases viral load [46]. The authors hypothesized that after a first infection, the memory T cells provide IFN-γ quickly following an additional exposure to another pathogen, similar to the study done with the H1N1 virus [46]. Additionally, IFN-γ mRNA was found to be upregulated during both JHMV and J2.2 infections [47]. It was shown to be essential for clearance of an attenuated strain of JHMV, J2.2, that leads to a persistent infection, in oligodendrocytes however, in the same study, it had little to no effect on viral clearance from neurons. This indicates a cell-specific role for IFN-γ immunity, which may affect the outcome of disease in the IFN-γ knockout mice [48].

Following our RNAseq experiments, we hypothesized that the increased IFN-γ was responsible for the restriction of MHV N1347A in the OB. This was largely based on a couple of pieces of evidence from previous work. First, we found that the KO of IFN-I signaling increases the replication of N1347A in BMDMs to near WT levels and also increased its pathogenesis. However, we did not detect much, if any, IFN-I in the OBs following JHMV infection. More recently we demonstrated that IFNγ pre-treatment induces PARP expression and has increased antiviral activity against a Mac1-deleted SARS-CoV-2 compared to WT virus [18]. In fact, several PARPs have been shown to be induced by both IFN-I and IFN-γ [49]. Thus, we hypothesized that in the absence of IFN-I, the production of IFN-γ would be responsible for increased PARP and ISG expression in the OB. However, we were surprised to find that the knockout of IFN-γ signaling did not enhance N1347A replication, but instead reduced the replication of both WT and N1347A viruses. It remains unclear what innate immune factors restrict N1347A in the OB, though one primary candidate is PARP14. PARP14 was one of the PARPs that was upregulated in the OB (Fig. 2D), and we have demonstrated that the knockout of PARP14 enhances N1347A replication in BMDMs [50]. However, PARP14 expression is not very robust in the OB, and thus other factors may play a role.

The more perplexing question is why is IFN-γ required for efficient JHMV replication in mice following intranasal infection **(Fig. 3)**? The addition of recombinant IFN-γ to BMDMs restricts virus replication **(Fig. 5C)**, and thus the requirement for IFNγ for JHMV replication during i.n. infection is likely due to the impact of IFN-γ on other cells in the brain. There was no significant difference in the levels of immune cells in WT vs IFN-γR^-/-^ mice, suggesting that IFN-γ may be more likely to impact non-hematopoetic cells in the brain that are susceptible to infection. IFN-γ is important for neuronal cell development [51], and the differentiation of neuronal precursor cells through cross talk with sonic hedgehog signaling [52]. Thus, it is possible that the lack of IFN-γ signaling from birth may impact olfactory neurons or other cells in the brain, such as astrocytes, that makes them less permissive to JHMV infection. It remains unclear what specific impacts IFN-γ has on cells in the OB that promote JHMV replication.

Finally, we were interested in the source of the IFN-γ that we observed at 5 dpi with JHMV. IFN-γ is primarily produced by CD8+ T cells and NK cells. However, other immune cells, such antigen-presenting cells, can produce IFN-γ as well. To determine which cell types were the predominant contributors of IFN-γ in the brain during JHMV infection, we quantified the percentage of immune cells expressing IFN-γ and the amount of IFN-γ production by different immune cells in the brains of infected animals using flow cytometry. Our results indicate that microglia constitute a significant proportion of immune cells in the brain **(Fig. 6)**. Further, the expression of IFN-γ in microglia was amongst the highest of immune cell populations we looked at (**Fig. 7B**). NK cells and CD4+ T cells produced comparably high levels of IFN-γ during infection as well (**Fig. 7B**). However, the proportion of CD4+ T cells and NK cells are less (**Fig. 6**), and in several mice these cells did not produce any IFN-γ (**Fig. 7A**). Taken together, we put forward that microglia are a major IFN-γ producing cell type following JHMV i.n. infection, though we can’t rule out contributions from T cells, NK cells, macrophages, and even non-hematopoetic cells. There are only limited reports demonstrating that microglia and other brain resident cells can produce IFN-γ, though evidence clearly demonstrates a role for microglia in the production of IFN-γ following Toxoplasma gondii infection [53–56]. This study provides additional evidence that microglia can be significant producers of IFN-γ following a virus infection in the brain. Future studies will be needed to further define how IFN-γ produced by microglia might impact both neuronal development and neurotropic virus infections.

## METHODS

### Cell Culture

HeLa cells expressing the MHV receptor carcinoembryonic antigen-related cell adhesion molecule 1 (CEACAM1) (HeLa-MHVR) were grown in Dulbecco’s Modified Eagle Medium (DMEM) supplemented with 10% fetal bovine serum (FBS), 100 U/ml penicillin and 100 mg/ml streptomycin, HEPES, sodium pyruvate, non-essential amino acids, and L-glutamine. Bone marrow-derived macrophages (BMDMs) sourced from WT, IFN-γ^-/-^ or IFN-γR^-/-^ mice were differentiated into M0 macrophages by incubating cells in Roswell Park Memorial Institute (RPMI) media supplemented with M-CSF (GenScript), 10% FBS, sodium pyruvate, 100 U/ml penicillin and 100 mg/ml streptomycin, and L-glutamine for six days. Then to differentiate into M2 macrophages, IL-4 (Peprotech Inc.) was added for 1 day. Cells were washed and replaced with fresh media every other day after the 4^th^ day.

### Mice

Pathogen-free C57BL/6 (B6) WT, IFN-γ, and IFN-γR knockout mice were originally purchased from Jackson Laboratories and mice were bred and maintained in the animal care facilities at the University of Kansas. Animal studies were approved by the University of Kansas Institutional Animal Care and Use Committees (IACUC) (permit #s 252-01 (ARF) and 278-01 (RCO) following guidelines set forth in the Guide for the Care and Use of Laboratory Animals.

### Virus Infection

Recombinant MHV-JHMV was previously described [20]. For intranasal infections, 5-8 week-old male and female mice were anesthetized with isoflurane and inoculated intranasally with 1×10^4^ PFU (unless otherwise stated) recombinant JHMV in a total volume of 12 μl DMEM. To obtain viral titers from infected animals, mice were sacrificed, and brain tissue was collected and homogenized in DMEM. Viral titers were determined by plaque assay using HeLa-MHVR cells.

### RNA Seq

RNA was isolated from C57BL6 mice either mock infected or infected i.n. with 1×10^4^ PFU N1347A virus using RNA purification columns (Qiagen). Library preparation was performed by the University of Kansas Genome Sequencing core facility, using the NEB Next RNA Library kit (NEB) with indexing. RNA-seq was performed using an Illumina NextSeq2000 high-output system with a paired-end reads of 50 bp each. Raw reads were processed with the nf-core RNA-seq pipeline v. 3.12.0 with Nextflow v. 23.04.3 [<u>10.5281/zenodo.7998767</u>]. Reads were trimmed of adapter sequences with TrimGalore v. 0.6.7 [<u>10.5281/zenodo.5127898</u>], assessed for batch effects and aligned to the C57BL6 mouse genome assembly version GRC m39 *Mus musculus* genome assembly (GCF_000001635.27) with the Ensembl version 81 annotation using STAR v 2.7.10a [57] and read count quantification was performed with RSEM v. 1.3.1 [58]. Differential gene expression between mock and N1347A infection treatments were estimated using DESeq2 v. 1.40.2 [59] and log_2_-fold change values were adjusted with the lfcShrink function specifying the apeglm method [60]. We classified those genes as differentially expressed that had Benjamini–Hochberg FDR-adjusted *P-*value (*P_FDR_*) < 0.01 and absolute log_2_ fold-change values > 2.

### Real-time qPCR Analysis

RNA was isolated from BMDMs using TRIzol (Invitrogen) and cDNA was prepared using MMLV-reverse transcriptase as per manufacturer’s instructions (Thermo Fisher Scientific). Quantitative real-time PCR (qRT-PCR) was performed on a QuantStudio3 real-time PCR system using PowerUp SYBR Green Master Mix (Thermo Fisher Scientific). Primers used for qPCR are listed in Table S2. Cycle threshold (C_T_) values were normalized to the housekeeping gene hypoxanthine phosphoribosyltransferase (HPRT) by the following equation: C_T_ = C_T(gene of interest)_ - C_T(HPRT)_. Results are shown as a ratio to HPRT calculated as 2^-ΔCT^.

### Immunohistochemistry (IHC)

At 5 dpi, mice were perfused intracardially with 4% formaldehyde (FA) diluted in 1X HBSS. After perfusion, each mouse brain was dissected and immersed in fresh 4% FA in individual tubes for post-fixation. The OB and forebrain regions were cut, placed on a freezing stage at −20°C and sectioned rostral-to-caudal at 30 µm in intervals (skip 60 µm between sections) using a sliding block microtome (American Optical Spencer 860 with Cryo-Histomat MK-2 controller). Four to six sections per group were moved individually on a 24-well plate using a camel hairbrush < 1.59 mm (Electron Microscopy Sciences, Cat. No. 65575-02), free-floating on HBSS/0.1% sucrose (HBSS/Su) at room temperature (RT) for rinsing. To perform single IHC, forebrain sections were permeabilized with HBSS/Su + 0.1% saponin (HBSS/Su/Sap; 2x, 5 min each), blocked with HBSS/Su/Sap + 0.1% Triton X-100 + 3% rabbit serum (1x, 1hr at RT), rinsed and incubated with the primary antibody mouse anti-N (1:5000) diluted in HBSS/Su/Sap + 3% rabbit serum O/N at 4 °C. The next day, sections were rinsed with HBSS/Su/Sap (3x, 5 min each) and incubated with the secondary antibody Alexa Fluor 594 rabbit anti-mouse (1:200) diluted in HBSS/Su/Sap (3 hrs at RT), rinsed with HBSS/Su/Sap (2x, 5 min each), HBSS/Su (1x, 5 min) and HBSS (1x, 5 min). Then Neurotrace^TM^ (1:25) and DAPI (10 µM) diluted in HBSS were added to stain for neurons and nuclear counterstain, respectively (1hr at RT), and then rinsed with HBSS (2x, 5 min each). Sections were carefully moved on a microscope slide using a camel hairbrush < 1.59 mm, mounted with Vectashield® Antifade mounting medium, and cover-slipped (22 x 50 cover glass; No. 1.5 thickness) for imaging.

### Image Acquisition and Quantification

Fluorescent images were acquired using a TCS SPE Laser Scanning Confocal Upright Microscope (Leica Microsystems, DM6-Q model), with the 405 nm and 561 nm laser lines, an Olympus 20X/0.75NA UPlanSApo infinity corrected, 8-bit spectral PMT detector and a Leica LAS X Imaging software (version 3.5.7.23225). Two to four images were taken per section. Anti-N + Alexa Fluor 594 signal was detected using 561 nm excitation (35% laser intensity), 600-630 nm emission range, 700 V PMT gain, and 0% offset. DAPI signal was detected using 405 nm excitation (6% laser intensity), 430-480 nm emission range, 680-700 V PMT gain, and 0% offset. Nissl signal was detected using 635 nm excitation (40% laser intensity, 650-700 nm emission range, 700-725 V PMT gain and 0% offset. Images were captured at 1024 x 1024-pixel resolution with a scan speed at 400, no bidirectional scanning, a zoom factor at 1.0, Pinhole 1.0 AU = 75.54 µm (550 µm x 550 µm image size; 537.63 nm x 537.63 nm pixel size; 2.057 µm optical section and 0.69 µm step size). Leica LAS X software_3D Viewer was used for post-processing to create figure plates, while raw data was exported as .tiff for relative fluorescent data analysis. In powerpoint, all images were set to 100% sharpness, 60% brightness, and 0% contrast. All the workflow design, sample preparation, processing and imaging was performed in the Microscopy and Analytical Imaging Resource Core Laboratory (RRID:SCR_021801) at The University of Kansas.

Quantification of N protein staining was automated using a custom macro written in FIJI (ImageJ v.1.54f). For each image, a median 3D filter was applied to single-channel Z-stacks (x=2, y=2, z=2) after which maximum intensity Z-projections were generated. Regions of Interest (ROIs) were generated following Medain 3D filtering (x=2, y=2, z=2) on duplicate single-channel Z-stacks followed by automated thresholding via the Triangle method. Resulting binary masks were applied to initial maximum Z-projections and fluorescence intensity of staining was measured within ROIs on the relevant channel. The total area of each ROI was also measured, and the Integrated Density (IntDen) displayed is a product of the mean fluorescence intensity and the ROI area.

### Flow Cytometry and Antibodies

Whole brains were excised and placed into RPMI with 10% FBS. Samples were processed with the gentleMACS Octo Dissociator (Miltenyi Biotec, Bergisch Gladbach, North Rhine-Westphalia, Germany) at 200 rpm without heat for 15 minutes. Following processing, dissociated samples were filtered through 70 uM filter to create a single-cell suspension in an ice-cold solution of 1 mL 10X PBS, 9 mL Percoll, and 10 mL RPMI. Tubes were centrifuged at 7900 rcf for 30 minutes, the myelin layer and media were then aspirated off, being careful not to disturb the pellet, leaving 10 mL of sample. The sample was centrifuged again at 1500 rcf for 10 minutes. Media was decanted and resuspended to desired concentration in RPMI with 10% FBS. Single cell suspension was then counted using the Vi-CELL BLU cell viability analyzer (Beckman Coulter, Brea, CA). Single cell suspensions were used for staining and flow cytometric analysis.

All flow cytometry was completed on a spectral cytometer the Cytek Aurora with a 5-laser system (405nm, 488nm, 640nm, 561nm, and 355nm). Single color antibody stain Positive and Negative Compensation Beads (BioLegend, San Diego, CA) were used for unmixing. Unmixed files were analyzed using FlowJo Software (BD Biosciences, San Diego, California). Antibodies used in various combinations (depending on experiment) are as follows: Ghost Viability Dye (Blue 516, BioLegend, 1:1000 dilution), CD4 (Super Bright 780, Invitrogen Thermo Fisher Scientific, 1:250, clone RM4-5), CD8 (APC-H7, BD Pharmingen,1:250, clone 53-6.7), NK1.1 (Super Bright 436, Invitrogen Thermo Fisher Scientific; or V450, BD Pharmingen, 1:250, clone PK136), CD45.R/B220 (APC-Cy5.5, Invitrogen Thermo Fisher Scientific, 1:250, Conjugate), F4/80 (Pacific Orange, Invitrogen Thermo Fisher Scientific, 1:250, Conjugate), CD3 (Super Bright 702, Invitrogen Thermo Fisher Scientific, 1:250, clone 17A2), CD45 (PerCP, BioLegend, 1:250, clone 30-F11), CD11c (Pe-Cy5.5, Invitrogen Thermo Fisher Scientific, 1:250, clone N418), CD11b (Super Bright 600, Invitrogen Thermo Fisher Scientific, 1:250, clone M1/70), Ly-6G (PE-eFluor 610, Invitrogen Thermo Fisher Scientific, 1:250, clone 1A8-Ly6g), CD19 (APC, BioLegend, 1:250, clone 6D5), CD44 (PE, eBioscience, 1:250, clone IM7), IFNγ (PE/Cy7, BioLegend, 1:100, clone XMG1.2). All surface markers were stained in 1% PBS, at 4°C, in dark. For intracellular cytokine staining of IFNγ, BD Cytofix and Permwash was used according to manufacturer’s instructions. If intracellular staining was being done, both surface and intracellular staining occurred following an incubation at 37°C with 5% CO_2_ for 4 hours with a BFA+RPMI.

### Statistics

All results are expressed as means ± standard errors of the means (SEM). Specific statistical tests used are indicated for each figure. P values of ≤0.05 were considered statistically significant (*, p≤0.05; **, p≤0.01; ***, p≤0.001; ****, p ≤0.0001; ns, not significant).

### Data and materials availability

All the RNAseq reads data are deposited in NCBI under the BioProject ID PRJNA1202193 and BioSample ID SAMN45953065 and SAMN45953641 and will be made public upon publication or March 31, 2025, whichever comes first. All other data will be made available with specific DOIs through the data repository FigShare upon publication.

## ACKNOWLEDGEMENTS

We thank members of the Davido laboratory at KU for valuable discussion. Research reported in this publication was made possible in part by the services of the KU Genome Sequencing Core which is supported by the National Institutes of General Medical Sciences of the National Institutes of Health under award number P30GM145499, the KU microscopy and analytical imaging (MAI) facility, the KU Center for Genomics, the K-INBRE Genomic Data Science Core supported by the IDeA Program which is supported by the National Institutes of General Medical Sciences of the National Institutes of Health under award number P20GM103418, and the KU Flow Cytometry Core which is supported by the National Institutes of General Medical Sciences of the National Institutes of Health under award number P20GM113117. We thank our funding from the NIH, an NIH Graduate Training grant, the University of Kansas College of Liberal Arts and Sciences, and the University of Kansas Madison and Lila Self graduate programs.

## Author contributions

Conceptualization: CMK, RCO, ARF

Data curation: CMK, MAP, SP, JJOC, RCO, ARF

Formal analysis: CMK, MAP, SP, JJOC, RCO, ARF

Funding acquisition: CMK, MAP, JJP, RCO, ARF

Methodology: CMK, MAP, SP, JJOC, JJP, RCO, ARF

Investigation: CMK, MAP, JP, JJP

Project administration: RCO, ARF

Resources: RCO, ARF

Visualization: CMK, MAP, SP, JJOC, RCO, ARF

Validation: CMK, MAP, SP, JJOC, RCO, ARF

Supervision: RCO, ARF

Writing—original draft: CMK, MAP, RCO, ARF

Writing—review & editing: CMK, MAP, SP, JJOC, JJP, RCO, ARF

A.R.F. was named as an inventor on a patent filed by the University of Kansas for a live-attenuated SARS-CoV-2 vaccine.

