## Supplemental Figures for "IFN-*γ* signaling is required for the efficient replication of murine hepatitis virus (MHV) strain JHM in the brains of infected mice"

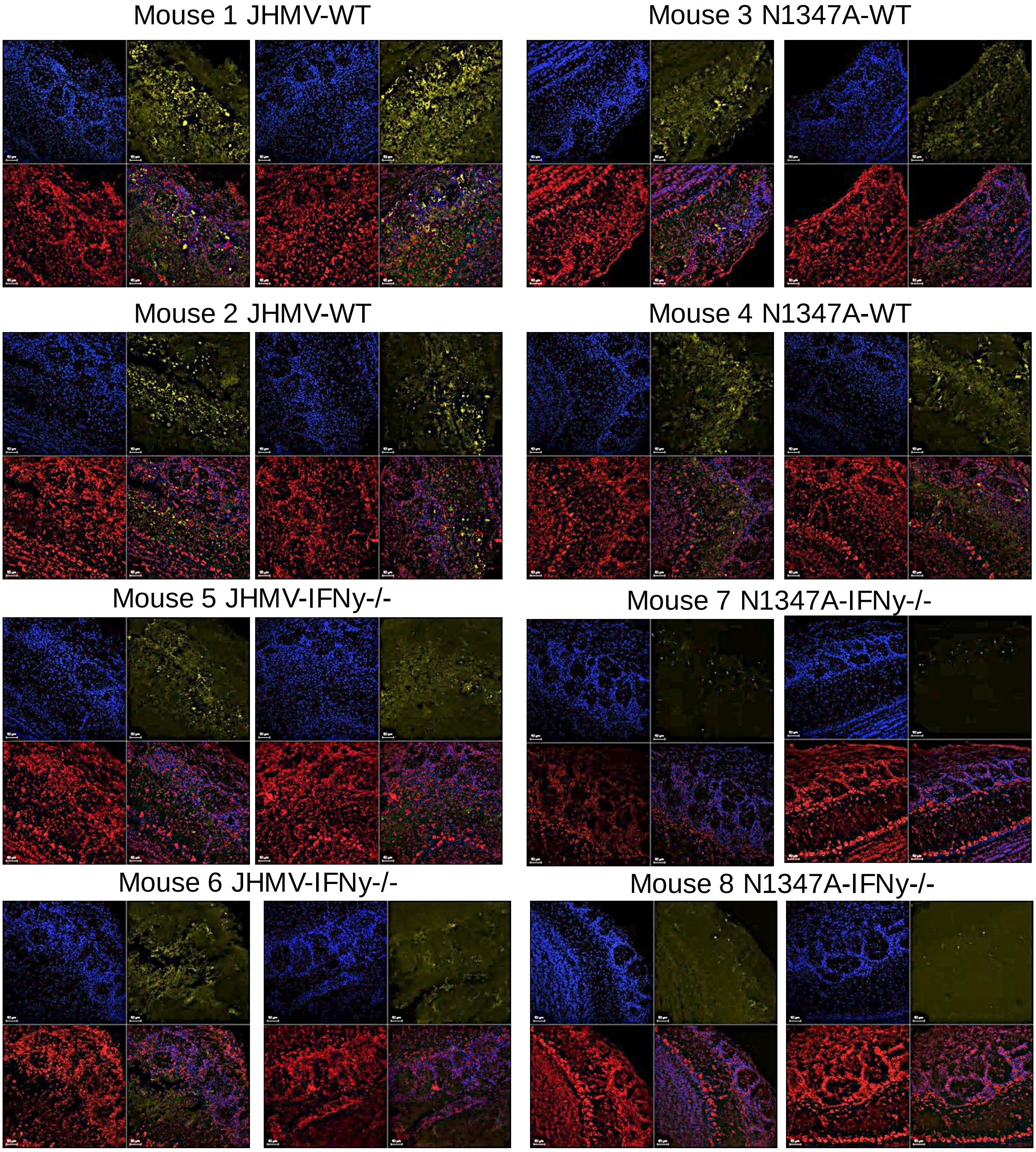


**Fig S1. JHMV replicates poorly in the brains of IFN-$\gamma$ null mice.** C57BL/6 WT and IFN-$\gamma^{-/-}$ mice were infected i.n. with $1{\times}10^4$ PFU WT or N1347A virus. At 5 dpi olfactory bulbs were fixed and sections were stained for MHV nucleocapsid (N) protein (yellow), DAPI (nuclei - blue) and Neurotrace™ (red) by IHC. n=2 mice for each group. Shown are two representative images for each mouse.

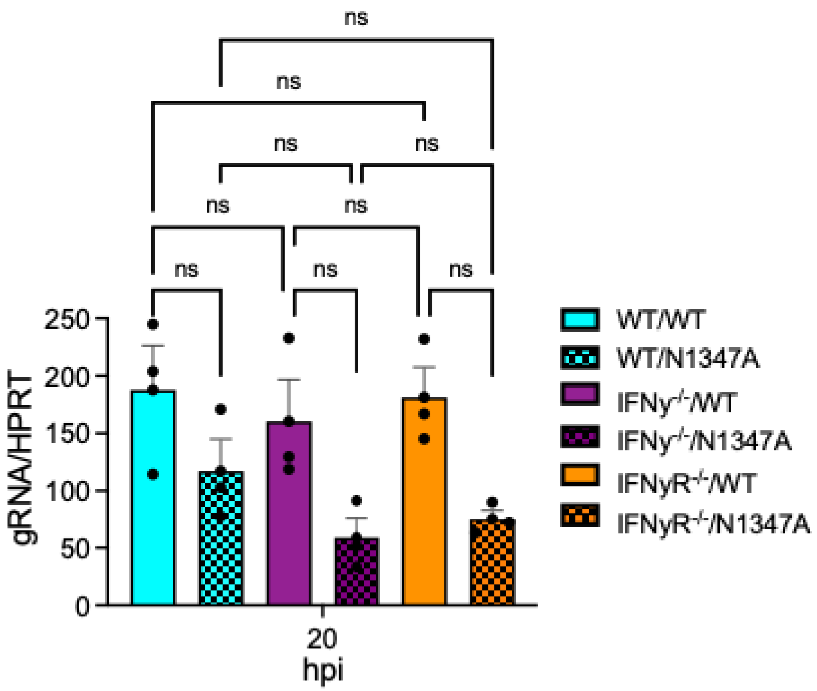


**Fig S2. The loss of IFN$\gamma$ in bone-marrow derived macrophages does not impact JHMV replication.** C57BL/6 WT, IFN$\gamma^{-/-}$, and IFN$\gamma$R$^{-/-}$ bone-marrow derived macrophages (BMDMs) were harvested from mice and differentiated into M2 macrophages as previously described. BMDMs were infected with WT and N1347A virus at an MOI of 0.1 and cells were collected at 20 hpi. RNA was harvested and JHMV genomic RNA was measured by qPCR. Data are from one experiment representative of two independent experiments. Statistics were determined by an ordinary one-way ANOVA.

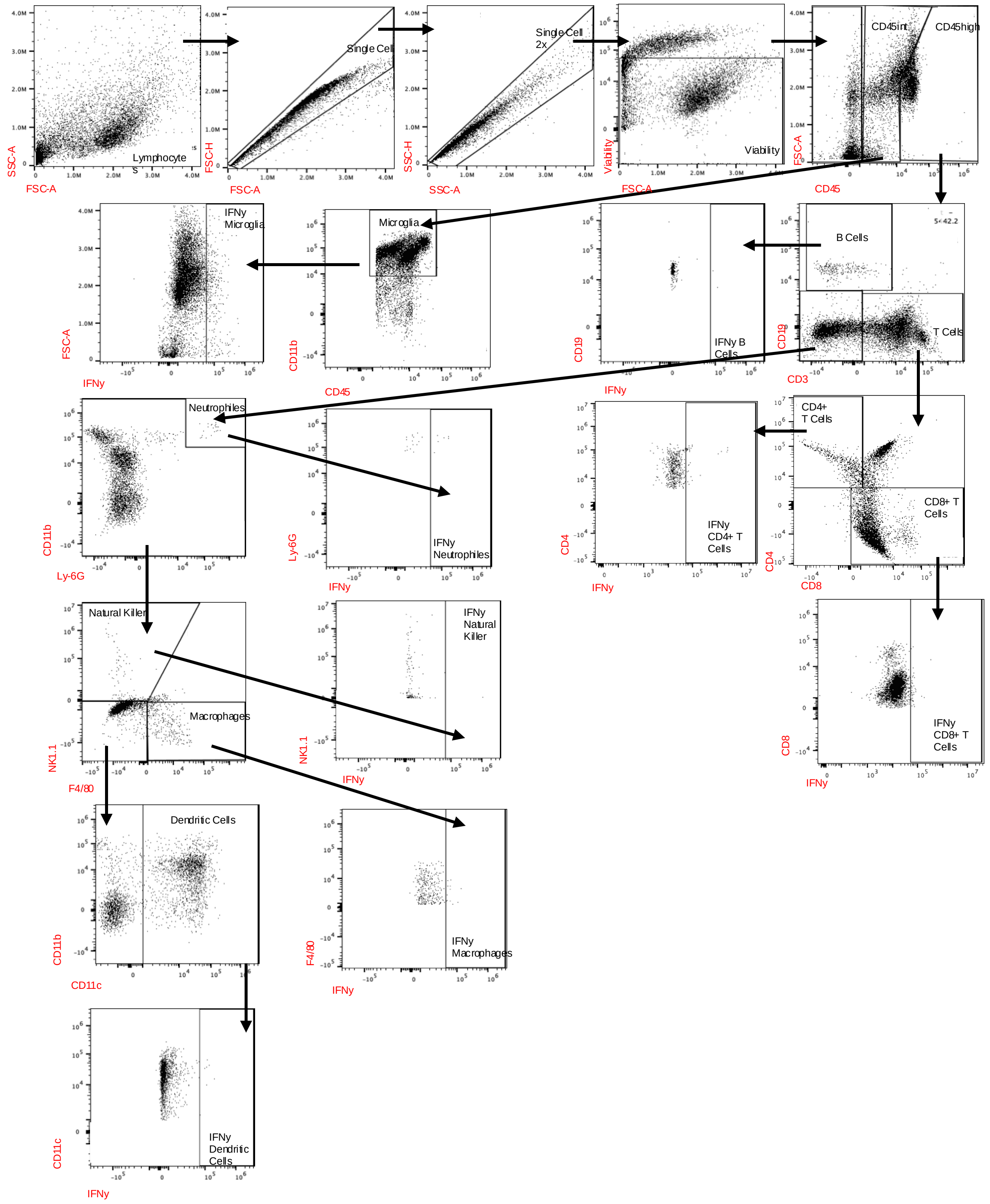


**Fig S3. Gating strategy for immune cells.**

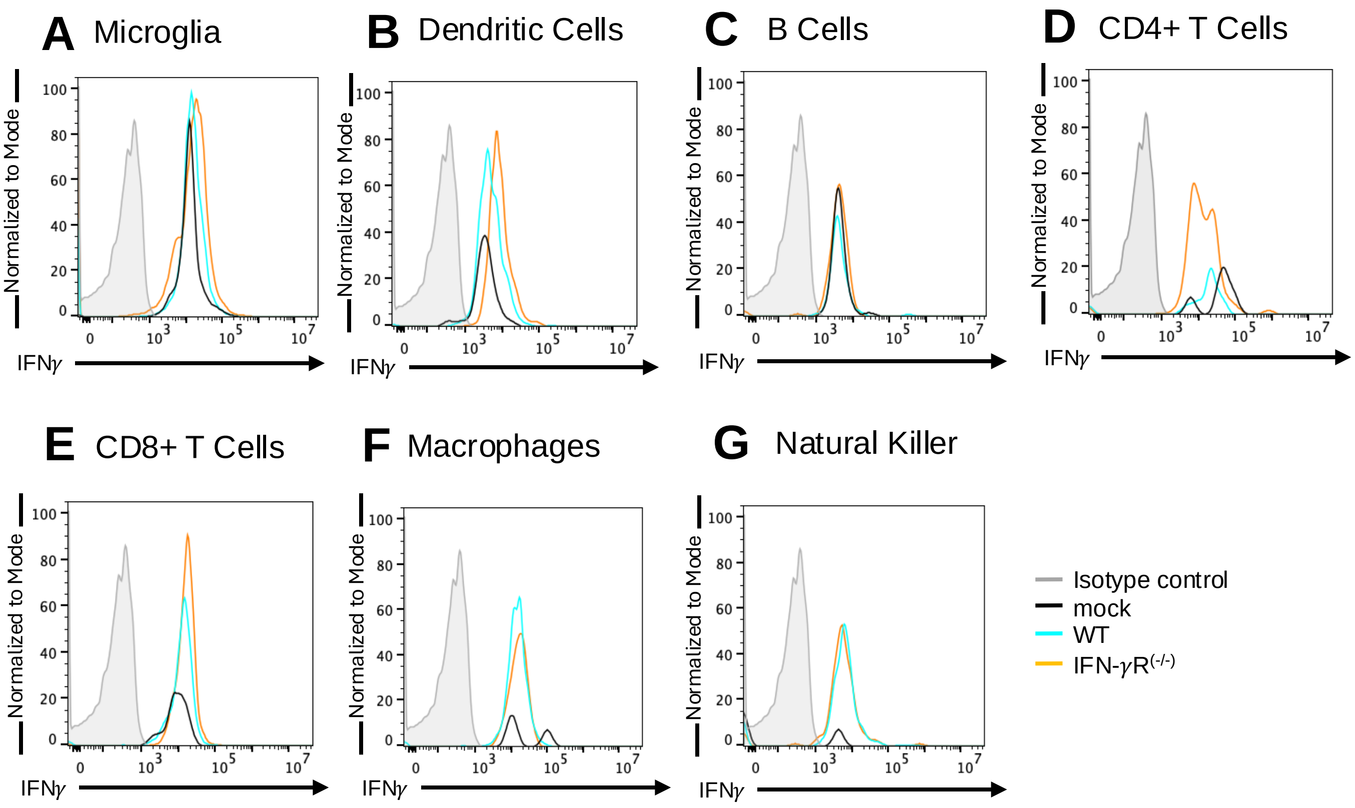


**Fig S4. Representative IFN-γ histograms in different cell populations.**
